# Astrocyte-specific inhibition of primary cilium functions improves cognitive impairment during neuroinflammation by suppressing A1 astrocyte differentiation

**DOI:** 10.1101/2023.10.01.560403

**Authors:** Nor Atiqah Muhamad, Shota Furukawa, Shunsuke Yuri, Michinori Toriyama, Kohei Masutani, Chuya Matsumoto, Seiya Itoh, Yuichiro Shinagawa, Ayako Isotani, Manami Toriyama, Hiroshi Itoh

**Author notes:** corresponding authors; Manami Toriyama and Hiroshi Itoh.

## Abstract

A1 astrocytes play a neurotoxic role in various neurodegenerative diseases. While inhibiting the differentiation of A1 astrocytes can slow disease progression, the mechanisms controlling A1 astrocyte differentiation are largely unknown. The primary cilium is a cellular organelle that receives extracellular signals and regulates cell proliferation, differentiation, and maturation. To elucidate the physiological function of the primary cilium in A1 astrocytes, we utilized primary astrocytes and an inflammation mouse model. We found that the length of the primary cilium was increased in astrocytes, and the inhibition of primary cilium formation inhibited their differentiation into A1 astrocytes. Since mice with systemic ciliogenesis defects exhibit embryonic lethality, the function of the primary cilium in adults has remained largely unclear. Therefore, we established conditional knockout (cKO) mice that specifically inhibit primary cilium function in astrocytes upon drug stimulation. In a neuroinflammation mouse model in which lipopolysaccharide (LPS) was intraperitoneally injected into wild-type mice, increases in A1 astrocyte number and primary cilium length were observed in the brain. In contrast, cKO mice exhibited a reduction in the proportions of A1 astrocytes and apoptotic cells in the brain. Additionally, the novel object recognition (NOR) score observed in the cKO mice was higher than that observed in the neuroinflammation model mice. These results suggest that the primary cilium in astrocytes is essential for A1 astrocyte differentiation, which leads to a decline brain function. We propose that regulating astrocyte-specific primary cilium signalling may be a novel strategy for the suppression of neuroinflammation.

## Background

The neural circuits in the CNS are composed of a range of cell types, including neurons and glial cells. Glial cells in the CNS are classified into three types: microglia, astrocytes, and oligodendrocytes (1). Astrocytes contribute to maintaining homeostasis, and the dysfunction of astrocytes is involved in various neurodegenerative diseases. For example, astrocytes regulate fluid and ion homeostasis, control blood flow and angiogenesis, protect neurons from excitotoxicity and cell death, promote synaptic formation, provide nutrients and energy metabolites to neurons, and are involved in bloodLbrain barrier (BBB) construction (2). Furthermore, astrocytes control excitatory synaptic transmission via astrocyte–neuron connections, influence microglial phenotypes, and induce phagocytosis via astrocyte–microglia crosstalk (3,4). Despite the well-known neuroprotective functions of astrocytes, a subtype that exerts neurotoxic effects, known as A1 astrocytes, has been identified (5). The differentiation of A1 astrocytes is induced by three cytokines secreted by microglia, namely, IL-1α, TNF-α, and C1q. An increased number of A1 astrocytes have been observed in the tissues of patients with neurodegenerative diseases, and they induce oligodendrocyte death by secreting saturated lipids (6). These reports suggest that inhibiting the differentiation or proliferation of A1 astrocytes may be a strategy for the prevention or treatment of neurodegenerative diseases. However, the detailed mechanisms controlling their differentiation are not yet clear.

The primary cilium is a nonmotile, single, microtubule-based, antenna-like organelle found on the surface of almost all cells types in vertebrates (7,8). The axoneme of the primary cilium protrudes from the cell surface, which enables extracellular signalling through a variety of ion channels and signalling receptors (9,10). Nearly 1000 cilium-related proteins have been recently identified in the primary cilium (11–14). Some of these proteins are localized at the ciliary membrane, whereas others are on the basal body and can be bidirectionally carried up and down the axoneme by the intraflagellar transport (IFT) complex. Some of these ciliary proteins and second messengers are involved in various signalling pathways. Moreover, the primary cilium is involved in Ca2+, Hedgehog (Hh), PDGFRIZ, Notch, TGF-β, mTOR, and other signalling pathways, which regulate cell proliferation, maturation, and differentiation (15–17).

Primary cilium assembly is tightly regulated. In vascular endothelial cells, intracellular and ciliary cAMP and cAMP-dependent protein kinase A (PKA) control cilium length and function (18). Several papers have reported that elevated ciliary cAMP levels increased primary cilium length (19–21). Mitogen-activated protein kinase (MAPK), protein phosphatase-1 (PP-1), and cofilin also control cilium length (18,21). These studies proposed that the molecular relationships between cilium function and length are not mutually exclusive (18). In addition, primary cilium length is regulated by inflammatory signalling. IL-1, nitric oxide and prostaglandin E2 (PGE2) elongate the primary cilium in articular chondrocytes via a PKA-dependent mechanism (22–25). Furthermore, recent evidence suggests a link between primary cilium defects and autoimmune and allergic diseases (26–29). These reports suggest that the primary cilium can regulate immune responses, although the role of the primary cilium in neuroinflammation remains largely unexplored. In this study, we investigated primary cilium function in A1 astrocytes. Our results suggest that the primary cilium plays a role in A1 astrocyte differentiation and propose that specific regulation of primary cilium signalling in astrocytes could be a therapeutic strategy for the treatment or prevention of neurodegenerative disorders.

## Methods

### Animal experiment

C57BL/6J mice were obtained from Japan SLC, Inc. Frozen embryos derived from B6.129P2-Ift88tm1Bky/J mice (#022409) were purchased from Jackson Laboratory (30). Frozen sperm derived from C57BL/6-Slc1a3(GLAST)<tm1(crePR)Ksak> (RBRC02350) mice were purchased from RIKEN BRC (31). Genetic modifications were verified by genotyping PCR using genomic DNA isolated from the mouse’s tail or ear tip. The following primers were utilized to identify the IFT88^+/+^ (WT, 365 bp), IFT88^flox/flox^ (∼410 bp), GLAST-Cre^-/-^ (WT, 120 bp), and GLAST-Cre^+/+^ (480 bp) alleles.

Fw-IFT88 floxed: GAC CAC CTT TTT AGC

CTC CTG Rv-IFT88 floxed: AGG GAA GGG ACT TAG GAA TGA

Fw-GLAST-Cre WT/MUT: TCTGGGAACAAGTGCAAGAC

Rv-GLAST-Cre WT: CTAGAGCCTGTTTTGCACGTT

Rv-GLAST-Cre MUT: TTGCTGCAATCTCTCCATCC

For the behavioural studies, mice aged 3 to 5 months were intraperitoneally (i.p.) injected with PBS and lipopolysaccharide (LPS, 1 mg/kg). LPS (Sigma, L2630) was dissolved in phosphate-buffered saline (PBS). To generate conditional knockout (cKO) mice in which exons 4-6 of *IFT88* were deleted (IFT88^flox^/^flox^ mice), GLAST-Cre^+/-^ mice were injected with 20 nmol/head RU486 diluted in 90% corn oil with 10% acetone twice a week for 2 weeks. Three days after the final administration of RU486, 1 mg/kg LPS was administered twice a week. Then, the mice were used in the behavioural studies. All experiments were performed in compliance with the Regulations and Laws of Animal Experimentation at the Nara Institute of Science and Technology (NAIST) and were approved by the Animal Experimental Committee of the NAIST (approval number: 1801). All efforts were made to minimize animal suffering.

### Behavioural studies

#### Novel object recognition (NOR) test

All mice were placed in an environment where the temperature was 23 to 24°C, the humidity was 50% to 70%, and the light/dark cycle was 12 h/12 h (lights at 8 am and turns off at 8 pm). The mice had free access to food and water. Two common objects were used a day before the test to acclimatize the mouse. On the training day, the common objects were presented to the mice for 10 minutes. After training, the mice were returned to their home cages and kept for 24 h. On the test day, a new object of the same material but a different shape was presented for 10 minutes, same as the original object. The mice were allowed to freely explore for 10 minutes, and the number of times the mice touched the object was calculated. The preference for the new object was calculated by the following formula and used as an index for long-term memory. New object preference = (The number of times touched to the new object)/(The number of times touched to the new object + original object) × 100. The objects were thoroughly cleaned with ethanol during each trial to remove the odour of other mice.

### Open field (OF) test

The behavioural tasks were performed in a soundproof room equipped with a behaviour analysis device, and the lighting of the soundproof room was set to 4 lux or less. The behaviour analysis device SCANET-40 (MELQUEST) was used for data acquisition, and the locomotor activity was determined by the corresponding software (MELQUEST, SCANET-40). In addition, mouse movement was recorded by a video camera. All mice were allowed one hour before the test to acclimate to the arena (45 cm × 45 cm) for 10 minutes. During the test, the mouse was placed in the central zone of the open field box. The mouse was permitted to explore the arena for 5 minutes. The entire arena was cleaned with 70% ethanol before the beginning of each test.

### Y-maze test

A mouse was placed in a Y-shaped apparatus with three arms at a 120 degree angle from each other and left for 7 minutes. The mouse was allowed to explore the apparatus freely. Squares, triangles, and cross marks were labelled on the wall behind the arm. The experiment was performed under 4 lux illumination. An arm entry was defined as the state in which the hind leg of the mouse passed over the centre of the arm, and the number of arm entries was recorded. The number of times the mouse entered the three different arms consecutively was recorded as the number of alternations, and the spontaneous alternation rate was calculated. Alternations = (Number of alternations)/((Total number of arm intrusion)-2) × 100. Data were obtained and used as an index of short-term memory.

### Astrocyte culture

A primary glial mixture was obtained from the cortices region of postnatal-7 (P7) pups as previously described (32) with minor modifications. After dissection of the brains, the meninges were removed, and the cortices were digested into small pieces in 4.7 ml of low calcium and high magnesium artificial cerebrospinal fluid (αCSF), 100 µl of 33.5 mg/ml hyaluronidase (Sigma, H3566), 100 µl of 5 mg/ml DNase (Merck, 10104159001) and 100 µl of 2.5% trypsin-EDTA (Life Technologies, 15090046) at 37°C for 15 min. Then, 50 µl of 70 mg/ml ovomucoid (Sigma, T9253) was added to stop the trypsin reaction, and the cortices were mechanically dissociated and resuspended in culture media (DMEM supplemented with 10% FBS) by repeated pipetting. The cell suspension was then filtered using a 70 µm nylon cell strainer (Falcon, 352350), and the cells were counted using a haematocytometer. Finally, 2 × 10^^6^ dissociated single cells were cultured as glial mixture cultures in a T75 flask and incubated at 37°C in a CO2 incubator. To obtain enriched astrocyte cultures, glial mixture cultures were shaken at 180 rpm for 30 minutes to remove microglia, and the flasks were shaken at 240 rpm for 6 h to remove oligodendrocyte precursor cells (OPCs). Next, the flask was vigorously shaken by hand for 1 minute to prevent OPC contamination. The old culture medium was removed, and the remaining confluent astrocytes were rinsed three times with PBS. New culture medium (DMEM supplemented with 10% FBS) was added, and the T75 flask was incubated at 37°C in a CO_2_ incubator. The culture medium was changed every 2-3 days.

### LPS and cytokine stimulation in primary cell cultures

LPS (Sigma, L2630) was diluted in D-PBS (Nacalai Tesque, 1424995). Mixed cortical glial cells were stimulated with 100 ng/ml LPS for 24 h, 48 h, and 72 h. Moreover, astrocyte cultures were treated with 3 ng/ml IL-1α (Sino Biological, 50114MNAE), 30 ng/ml TNF-α (PeproTech, 30001A), and 400 ng/ml C1q (Merck, 204876) for 24 h.

### RNA interference

*IFT88*-targeted small interfering RNA (siRNA) was purchased from Life Technologies (P/N4390815) with the following 5’ to 3’ sequences: sense: GGACUUAACCUACUCCGUUtt and antisense: AACGGAGUAGGUUAAGUCCaa). Based on the manufacturer’s recommended protocol, knockdown was performed for 48 h using Lipofectamine RNAiMAX (Invitrogen, P/N56532). Silencer® Negative Control siRNA #1 (Ambion, AM4611) was used as the control. Two days after transfection, the culture medium was changed, and the cells were stimulated with 3 ng/ml IL-1α, 30 ng/ml TNF-α, and 400 ng/ml C1q for 24 h.

### Immunofluorescence imaging

Following fixation in 4% paraformaldehyde (Nacalai Tesque, 2612654) in PBS at 37°C for 20 minutes, coverslip cultures were washed three times with PBS. Samples were permeabilized and blocked with blocking buffer containing 0.1% Triton X-100 and 5% foetal calf serum in PBS for 30 minutes. Then, the samples were incubated with primary antibodies (Table 1) diluted with blocking buffer and incubated overnight at 4°C. The samples were washed three times with PBS, incubated with secondary antibodies and Hoechst 33342 to detect nuclei (Table 1), and diluted with blocking buffer for 1 h at room temperature with light protection. The samples were washed with PBS three times every 10 minutes, and coverslip glass was mounted using Mountant PermaFluor (Thermo, TA030FM). Fluorescence microscopes (Axio Observer Z1(Carl Zeiss); FV1200 confocal microscope (Olympus)) were used for slide observation.

**Table 1.** Primary and secondary antibodies used for immunofluorescence imaging.

| Primary antibodies | Concentration | Source (Catalogue Number) |
| --- | --- | --- |
| Mouse anti-ARL13B monoclonal antibody | 1:100 (brain), 1:1000 (cell) | Abcam (ab136648) |
| Rabbit anti-ARL13B polyclonal antibody | 1:500 (brain), 1:1000 | Proteintech (17711-1-AP) |
| Rabbit anti-GFAP polyclonal antibody | 1:1000 | Dako (Z0034) |
| Mouse anti-GFAP monoclonal antibody | 1:100 | Sigma-Aldrich (MAB360) |
| Rat anti-C3 monoclonal antibody | 1:50 (brain), 1:500 (cell) | Santa Cruz (sc58926) |
| Rabbit anti-Cre recombinase | 1:50 | Cell Signaling Technology (15036T) |
| Secondary antibodies and reagent | Concentration | Source (Catalogue Number) |
| Hoechst 33342 | 1:1000 | Thermo (H1399) |
| Alexa Fluor™ 488 anti-mouse IgG | 1:1000 | Life Technologies (A21202) |
| Alexa Fluor™ 594 anti-mouse IgG | 1:1000 | Invitrogen (A11032) |
| Alexa Fluor™ 488 anti-rabbit IgG | 1:1000 | Invitrogen (A48282) |
| Alexa Fluor™ 594 anti-rabbit IgG | 1:1000 | Life Technologies (A21207) |
| Alexa Fluor™ 488 anti-rat IgG | 1:1000 | Life Technologies (A21208) |

Perfusion and fixation was performed using PBS and 4% PFA in PBS after behavioural testing. After perfusion and fixation, an incision was made in the parietal region of the mouse, and the brain was extracted. The excised brain was fixed again with 4% PFA in PBS at 4°C for 24 h and then dehydrated in a sucrose-PBS gradient (10%, 20%, 30% w/v), and the brain was embedded in OCT compound (Sakura Finetech Japan, Inc., 45833) and frozen at -80°C. Coronal sections with a thickness of 10 μm were prepared using a cryostat. The cryosections prepared above were fixed in 4% PFA in PBS for 10 minutes and then prechilled to -30°C. Next, cryosections were dehydrated and defatted in ice-cold acetone for 10 minutes, followed by washing with PBS three times. Afterwards, the samples were immersed in 1 mM EDTA in PBS (pH 8.0) at 70°C for 20 minutes for antigen retrieval. Cryosections were permeabilized with 0.2% Triton-X100 in PBS at room temperature for 10 minutes and washed with PBS three times. Then, the cells were blocked with 5% BSA, 0.1 M glycine, 10% FcR blocking reagent (Miltenyi Biotec, #130-092-575) and 0.1% Tween 20 in PBS for 1 h at room temperature. The primary antibodies (Table 1) were diluted with 5% BSA and 0.1% Tween 20 in PBS and incubated at 4°C overnight. Washing with PBS was performed three times every 10 minutes, and the secondary antibodies (Table 1) were diluted with 5% BSA and 0.1% Tween 20 in PBS containing Hoechst 33342 for 1 h at room temperature while shielded from light. Again, washing with PBS was performed three times every 10 minutes, and cover glass samples were mounted using mounting medium. Observations were made using a fluorescence microscope (Axio Observer Z1(Carl Zeiss); FV1200 confocal microscope (Olympus)).

## Results

### A1 astrocyte differentiation increases primary cilium length

While it has been reported that brain astrocytes can assemble the primary cilium, its physiological functions remain largely unexplored. Additionally, although there have been numerous suggestions regarding the role of the primary cilium in immune response regulation, detailed analyses are lacking. To investigate the physiological functions of the primary cilium in neurotoxic A1 astrocytes, we isolated glial cells from the brains of postnatal day 7 (P7) mice and induced A1 astrocyte differentiation by stimulating them with LPS (Fig 1A) (5). Subsequently, we observed the expression of the A1 astrocyte marker C3 and the primary cilium in cultured astrocytes. Stimulation with LPS in mixed glial cell cultures upregulated C3 expression in GFAP-positive cells (Fig S1A, B). Under these conditions, more than 90% of the GFAP-positive cells formed primary cilia, and the primary cilium formation rate remained unchanged following LPS stimulation (Fig 1B, C). However, LPS stimulation in mixed glial cell cultures significantly increased the length of the primary cilium in GFAP-positive astrocytes (Fig 1B, D).

**Figure 1.**
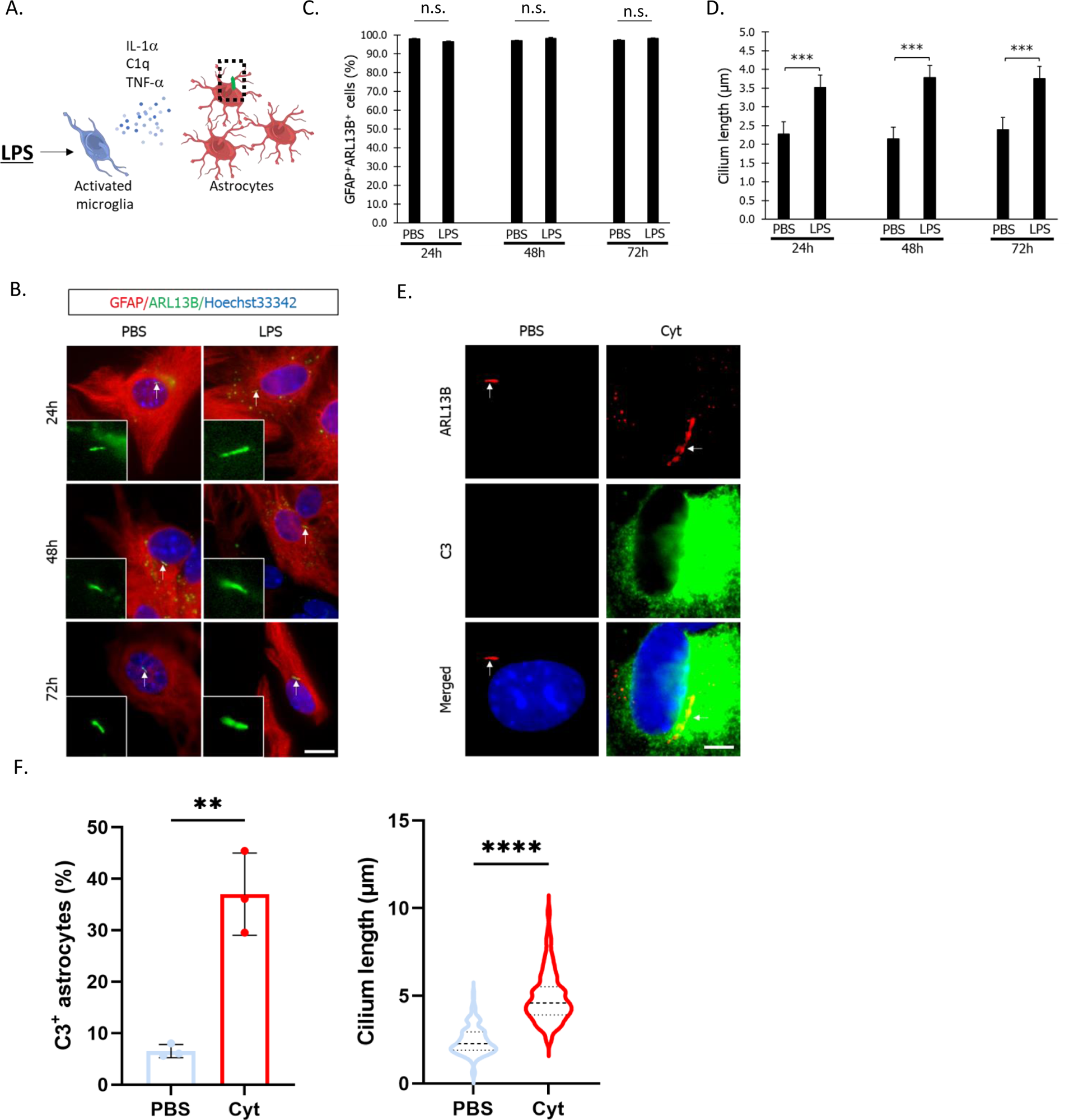
A1 astrocyte differentiation increases primary cilium length. (A) Schematic illustration of A1 astrocyte differentiation. Microglial activation by LPS stimulation upregulates the expression of IL-1α, C1q, and TNF-α, facilitating the differentiation of A1 astrocytes (5). (B) Representative immunostaining image of Arl13B (primary cilium, green) in GFAP-positive astrocytes (red). Mixed cortical glial cells were treated with 100 ng/ml LPS for 24 h, 48 h, and 72 h. The arrow indicates the primary cilium. Nuclei were stained with Hoechst 33342 (blue). Scale bar; 20 μm. (C) Percentage of astrocytes with a primary cilium. Primary astrocytes were stimulated with PBS or 100 ng/ml LPS for 24, 48, 72 h. n=3 independent experiments. The Mann‒Whitney U test was performed. ns, nonsignificant. (D) The astrocytic primary cilium length shown in Figure 1B was measured. n=150, 3 independent experiments were performed. ***, P <0.001 (Mann‒Whitney U test). (E) Representative image of Arl13B (red) and C3 (green) and nuclei (Hoechst 33342, blue) in astrocytes. Astrocytes were derived from glial mixture culture and stimulated with PBS or a cytokine mixture (3 ng/mL IL-1α, 30 ng/ml TNFα, 400 ng/ml C1q) for 24 h. The arrow indicates the primary cilium. Scale bar; 5 μm. (F) (left) Percentage of C3-positive astrocytes shown in Figure 1 E. Three independent experiments were performed. **, P <0.01, (Mann‒Whitney U test). (Right) Astrocytic cilia length is shown in the graph (n=150, three independent experiments). ****, P <0.0001. (Mann‒Whitney U test).

It has been reported that astrocytes differentiate into A1 astrocytes in response to three cytokines secreted by microglia, IL-1IZ, TNF-IZ, and C1q (5). Therefore, we stimulated astrocytes from glial cultures with these cytokines and observed the primary cilium after stimulation. As expected from the results obtained using mixed glial cell cultures, the cytokine stimulation of astrocytes increased the proportion of C3-positive cells, accompanied by increased primary cilium length (Fig 1E, F). To further investigate the correlation between A1 astrocyte differentiation and primary cilium length, we examined the time it took for the expression level of C3 and cilium length to increase following cytokine stimulation. It was suggested that both C3-positive cell numbers and primary cilium length gradually increased after cytokine stimulation, starting from 1 h poststimulation (Fig S1C, D). These results suggest that LPS activates microglia, promoting cytokine production and A1 astrocyte differentiation, accompanied by primary cilium elongation.

### Inflammation model mice exhibit an increase in A1 astrocyte numbers, elongation of the primary cilium, and a decline in cognitive function

To observe the changes in A1 astrocytes numbers and primary cilium length in the brains of adult mice, we established an inflammation model by intraperitoneally (i.p.) administering LPS. In the inflammation model mice, the number of C3-positive astrocytes were increased compared that in the control group treated with PBS, which was in accordance with previous findings (5) (Fig S2A, B). When observing the primary cilium, we found that the length of primary cilia in both GFAP-positive cells and C3-positive cells was significantly increased compared to that in the control group (Fig 2A, B, S2C, D). The primary cilium formation rate remained unchanged in both GFAP-positive and GFAP-negative cells (Fig 2B, S2D). These results suggest that inflammation induced by i.p. LPS administration elongates the primary cilium in astrocytes in the brain. It has been reported that inflammation model mice exhibit a decline in cognitive function (33–35). Therefore, we investigated mouse cognitive function using the novel object recognition (NOR) test for evaluating long-term memory, the open field (OF) test for determining locomotor activity, and the Y-maze test for assessing short-term memory (Fig 2C). In the inflammation model mice, the preference for the novel object was significantly lower than that of the control mice, whereas no significant differences were observed in the OF test or Y-maze test (Fig 2D). These results suggest that in inflammation model mice, the elongation of the primary cilium in A1 astrocytes is associated with a decline in long-term memory.

**Figure 2.**
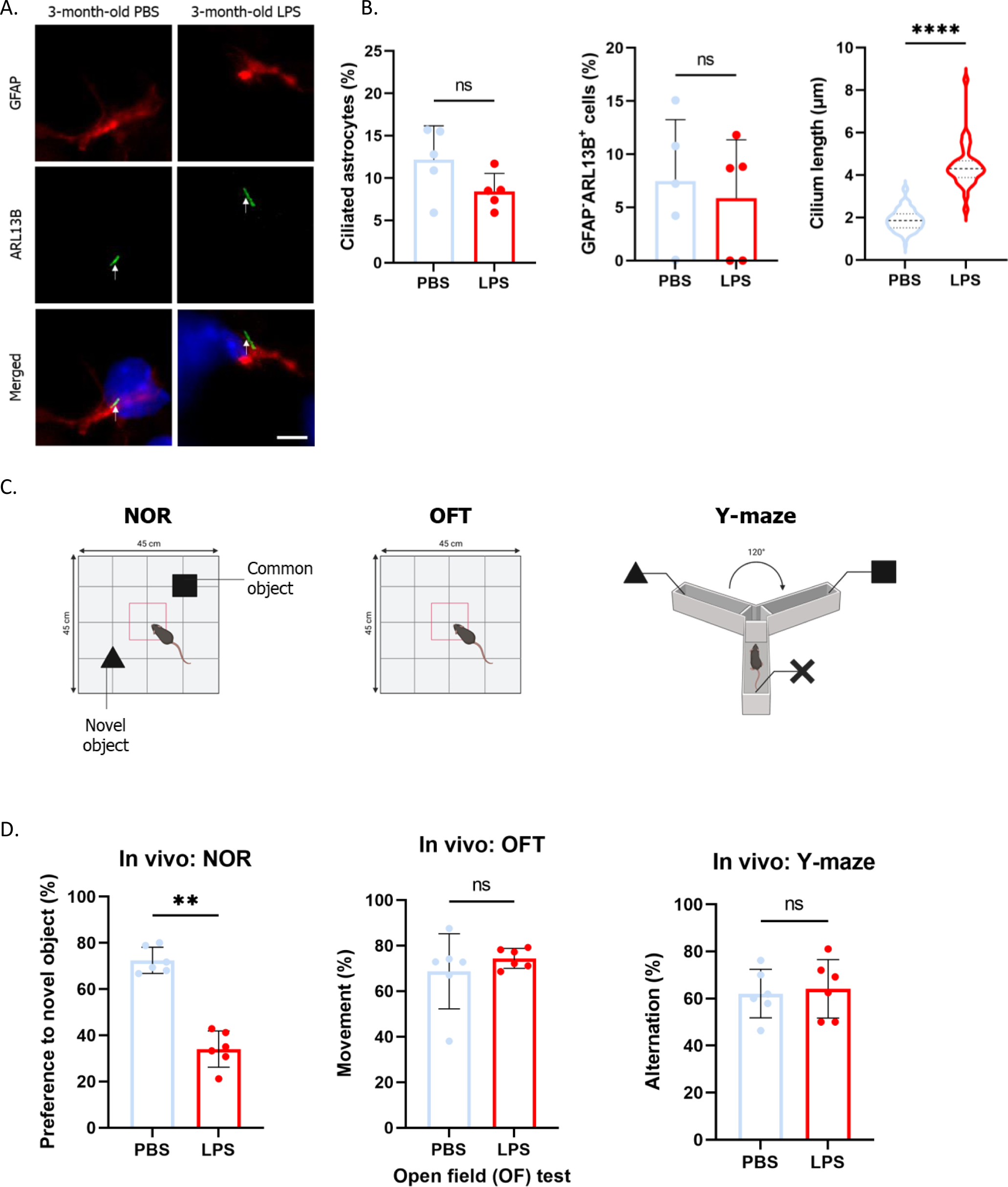
LPS injection elongates the astrocytic primary cilium and reduces cognitive function in mice. (A) Representative image of GFAP (astrocytes, red), Arl13B (primary cilium, green), and nuclei (blue) in the mouse brain. Three-month-old mice were i.p. injected with 1 mg/kg LPS twice a week for 6 weeks. Scale bar; 10 μm. PBS-treated mice; n=5. LPS-treated mice; n=5. (B) The percentage of ciliated GFAP-positive astrocytes (left), percentage of ciliated GFAP-negative cells (middle), and astrocytic primary cilium length (right) shown in Figure 2A are shown in the graph. ****, P <0.0001, ns, nonsignificant. (Mann‒Whitney U test). (C) Illustration of the mouse behavioural test. The novel object recognition (NOR) test (left), open field (OF) test (middle) and Y-maze test (right) were performed. Figures were created with BioRender.com. (D) Results of behavioural (object recognition) studies in 3-month-old PBS- and LPS-injected C57BL/6J mice. PBS-treated mice; n=6. LPS-treated mice; n=6. **, P <0.01, ns, nonsignificant. (Mann‒Whitney U test).

### The specific inhibition of primary cilium formation in astrocytes suppresses A1 astrocyte differentiation and ameliorates LPS-induced cognitive impairment

IFT88 is essential for primary cilium formation and maintenance, and its downregulation has been known to inhibit primary cilium formation (36,37). To investigate the physiological function of the primary cilium in A1 astrocyte differentiation, we transfected siRNA targeting *IFT88* into cultured astrocytes and quantified the number of C3-positive astrocytes after stimulation with cytokines. Knockdown of *IFT88* suppressed primary cilium formation in astrocytes (Fig S3A-C). Furthermore, *IFT88* knockdown significantly decreased the percentage of C3-positive astrocytes, although the cells were stimulated with cytokines (Fig S3D).

Since our results suggested that the inhibition of primary cilium formation in astrocytes suppresses A1 astrocyte differentiation, we established conditional knockout (cKO) mice in which exons 4-6 of the *IFT88* gene were specifically deleted in astrocytes. Then, RU486 was administered and the phenotype of the mice was observed (30,31,38). We crossed *IFT88*^flox^ mice with mice expressing CrePR under the control of astrocyte-specific *GLAST* promoter (*GLAST-CrePR*) activation and intraperitoneally administered RU486 to 8-week-old offspring (Fig S4A). After RU486 administration, we intraperitoneally injected LPS, performed an OR test, and determined C3 expression in astrocytes (Fig 3A). To initially confirm the efficiency of *IFT88* reduction, we measured the primary cilium formation rate in mouse brain astrocytes. We first tried to detect IFT88-positive primary cilia in the brain; however, the signal was extremely low even in wild-type mice, so we calculated the primary cilium formation rate by detecting Arl13B, which is commonly used as a primary cilium marker. In RU486-administered *IFT88*^flox/flox^ *GLAST-CrePR* (cKO) mouse brains, a reduced primary cilium formation rate was observed in GFAP-positive cells (Fig 3B, C). In contrast, there was no significant difference in the primary cilium formation rate in GFAP-negative cells between *IFT88*^flox/+^; *GLAST-CrePR* (ctrl) and cKO mice (Fig 3C).

**Figure 3.**
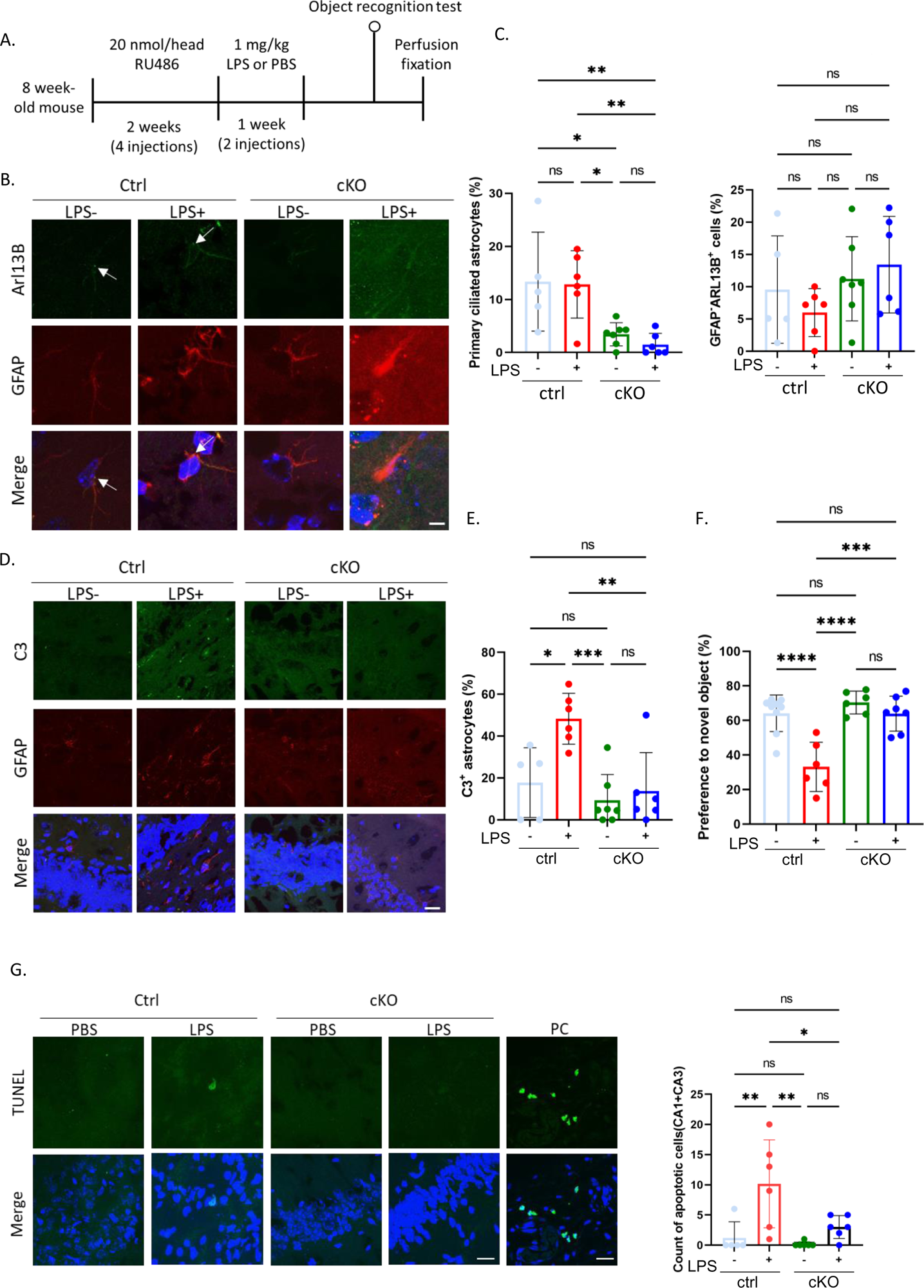
Astrocyte-specific *IFT88* gene knockout attenuates cognitive decline induced by LPS injection. (A) Timeline of experiments. Eight-week-old mice (IFT88^flox/flox^; GLAST-CrePR^-/-^ (ctrl) or IFT88^flox/flox^; GLAST-CrePR^+/-^ (cKO)) were i.p. injected with 20 nmol/head RU486 twice a week for 2 weeks. Three days after the final injection of RU486, the mice were i.p. injected with 1 mg/kg LPS twice a week. A separate group was also prepared and an equal volume of PBS was administered. Two days after the second LPS injection, an NOR test was performed. (B) Representative image of Arl13B (green), GFAP (red) and nuclei (Hoechst 33342, blue) in the mouse brain. Scale bar; 5 μm. The arrow indicates the primary cilium. PBS administration to ctrl mice; n=5. LPS administration to ctrl mice; n=6. PBS administration to cKO mice; n=7. LPS administration to cKO mice; n=6. (C) Percentage of ciliated astrocytes in the brain (left) and percentage of ciliated GFAP-negative cells (right) shown in Figure 3B. *, P <0.05, **, P <0.01, ns, nonsignificant. (Tukey’s multiple comparison). (D) Representative image of C3 (green), GFAP (red) and nuclei (Hoechst 33342, blue) in the mouse hippocampal area. Scale bar; 20 μm. PBS administration to ctrl mice; n=5. LPS administration to ctrl mice; n=6. PBS administration to cKO mice; n=7. LPS administration to cKO mice; n=6. (E) Percentage of C3-positive astrocytes shown in Figure 3D. PBS-treated ctrl mice; n=9. LPS-treated ctrl mice; n=6. PBS-treated cKO mice; n=6. LPS-treated cKO mice; n=7. *, P <0.05, **, P <0.01, ***, P <0.001, ns, nonsignificant. (Tukey’s multiple comparison). (F) Results of ORT studies in mice. PBS-treated ctrl mice; n=9. LPS-treated ctrl mice; n=6. PBS-treated cKO mice; n=6. LPS-treated cKO mice; n=7. ***, P <0.001, ****, P <0.0001, ns, nonsignificant. (Tukey’s multiple comparison). (G) A representative image of early apoptotic cells detected by the TUNEL method in mouse hippocampal CA1/CA3 regions (left). Scale bar; 20 μm. PBS-treated ctrl mice; n=9. LPS-treated ctrl mice; n=6. PBS-treated cKO mice; n=6. LPS-treated cKO mice; n=7. Positive control (PC) slides were used as staining controls. The number of TUNEL-positive cells is shown in the graph (right). *, P <0.05, **, P <0.01, ns, nonsignificant. (Tukey’s multiple comparison).

These results confirm the astrocyte-specific reduction in the primary cilium in cKO mice. Subsequently, we examined C3 expression in these mice after LPS administration. In control mice, an increase in the number of C3-positive astrocytes was observed upon LPS administration, while no significant increase in C3-positive astrocyte numbers was observed in cKO mice even after LPS administration (Fig 3D, E). We also examined primary cilium disruption in cultured astrocytes. The inhibition of primary cilium formation was not observed in control mouse-derived or C57BL/6J (wild-type B6J) mouse-derived cultured astrocytes, indicating that it was a feature of only cKO mouse-derived astrocytes treated with RU486 (Fig S4B, C, E, F). Furthermore, the increase in C3-positive astrocyte numbers induced by LPS stimulation was not observed in cKO mouse-derived astrocytes (Fig S4G, H).

Furthermore, cognitive tests were conducted on control and cKO mice.

In control mice, the preference for the novel object was reduced upon LPS administration, whereas a decline in the preference for the novel object was not observed even after LPS administration in cKO mice (Fig 3F). Compared to wild-type, mice cKO mice without RU486 administration did not show any differences in cognitive function (Fig S4D).

It has been reported that A1 astrocytes induce oligodendrocyte death by secreting saturated lipids (5,6). To examine whether A1 astrocytes induce cell death in the brain, we detected apoptotic cells upon LPS administration using the TdT-mediated biotin-dUTP nick end labelling (TUNEL) method. In control mice, LPS administration led to an increase in the proportion of TUNEL-positive cells near the hippocampal CA1/CA3 regions, whereas no significant increase in the proportion of TUNEL-positive cells was observed in cKO mice (Fig 3G). These results suggest that the functional inhibition of the primary cilium in astrocytes suppressed A1 astrocyte differentiation and attenuated cognitive decline by suppressing cell death in the brain.

## Discussion

In this study, we showed that the length of the primary cilium was elongated in A1 astrocytes compared with unstimulated astrocytes. Additionally, astrocyte-specific inhibition of primary cilium formation suppressed A1 astrocyte differentiation and attenuated cognitive decline. These results suggest the importance of primary cilium signalling in A1 astrocyte differentiation; however, the physiological relevance of primary cilium elongation in A1 astrocytes remains unclear. Primary cilium elongation could potentially alter the density of signalling molecules present in the primary cilium, thereby affecting downstream signalling (21). Interestingly, it has been suggested that not only the regulation of primary cilium formation but also the modulation of cilium length, formation rate, and receptor localization changes under different physiological conditions in the brain (39). Cilium length regulation may be involved in the control of A1 astrocyte differentiation by modulating immune signalling pathways.

Stimulation with TNFα, C1q, and IL-1α elongated the primary cilium in astrocytes (Fig 1E, F). Previous reports have indicated that both IL-1α and IL-1β elongate the primary cilium in chondrocytes by upregulating the expression of inflammatory mediators such as COX2 and iNOS, and this effect is inhibited by the knockout of *IFT88* (22–24). Moreover, COX2 was upregulated upon IL-1β stimulation in cells in which the expression of *Kif3A* and *TTBK2*, which are also essential for primary cilium formation and maintenance similar to IFT88, was inhibited (25). These findings suggest that there is a primary cilium-independent mechanism that regulates the expression of inflammatory mediators or that the inflammatory signalling within cilia is intricately controlled by multiple molecules (25). Further research is required to understand the mechanisms by which the primary cilium regulates inflammatory signalling.

Numerous studies have suggested the importance of the primary cilium in neuronal cells (40). Intraperitoneal LPS administration in mice shortens primary cilium length in hippocampal neurons (41). Furthermore, the neuron-specific inhibition of primary cilium formation showed no impact on short-term memory but resulted in impaired long-term memory retention (42). The primary cilium has also been suggested to form an axon-cilium synapse in the brain (43). These findings strongly suggest that the primary cilium plays a crucial role in maintaining homeostasis in neuronal cells. Recently, it has become a concern that, even long after recovery from COVID-19, brain structural atrophy and cognitive decline are still observed (44). Systemic inflammation induced by the activation of Toll-like receptors (TLRs) and inflammatory mediators such as TNFα has been suggested to cause cognitive decline and accelerate the progression of Alzheimer’s disease (45,46). Considering the findings of this study and the importance of the primary cilium in neuronal cells, it is suggested that temporally-dependent and astrocyte-specific suppression of the primary cilium, especially during inflammation induction, may help maintain brain homeostasis. In the future, elucidating the ciliary signalling pathways within the primary cilium of A1 astrocytes will contribute to the development of therapeutics for neuroinflammation.

## Conclusion

Astrocyte-specific suppression of the primary cilium suppressed A1 astrocyte differentiation induced by LPS administration and improved cognitive function. Identification of important primary cilium signalling pathways in A1 astrocytes will contribute to the development of novel therapeutics for neuroinflammation.

## Abbreviations

LPS: lipopolysaccharide
IFT: intraflagellar transport
PGE2: prostaglandin E2
IL-1: interleukin-1
TNFα: tumour necrosis factor α
PDGFRα: platelet-derived growth factor receptor α
TGFβ: transforming growth factor β
mTOR: mammalian target of rapamycin
cAMP: cyclic adenosine monophosphate
GFAP: glial fibrillary acidic protein
GLAST: L-glutamate/L-aspartate transporter
iNOS: inducible nitric oxide synthase
COX2: cyclooxygenase 2

## Author Contributions

NA M, S F., S Y., Mi T., C M., S I., Y S., and Ma T. conducted the experiments. Ma T., An I., and H I conceived and designed the experiments and wrote the manuscript. NA M., Ma T., and Mi T. contributed to the writing and editing of the manuscript.

## Acknowledgements

We greatly appreciate Dr. Kenji Sakimura at Niigata University for providing the C57BL/6-Slc1a3(GLAST)<tm1(crePR)Ksak> mice. We very much thank Chie Murayama for her excellent technical assistance.

---

**Figure S1.**
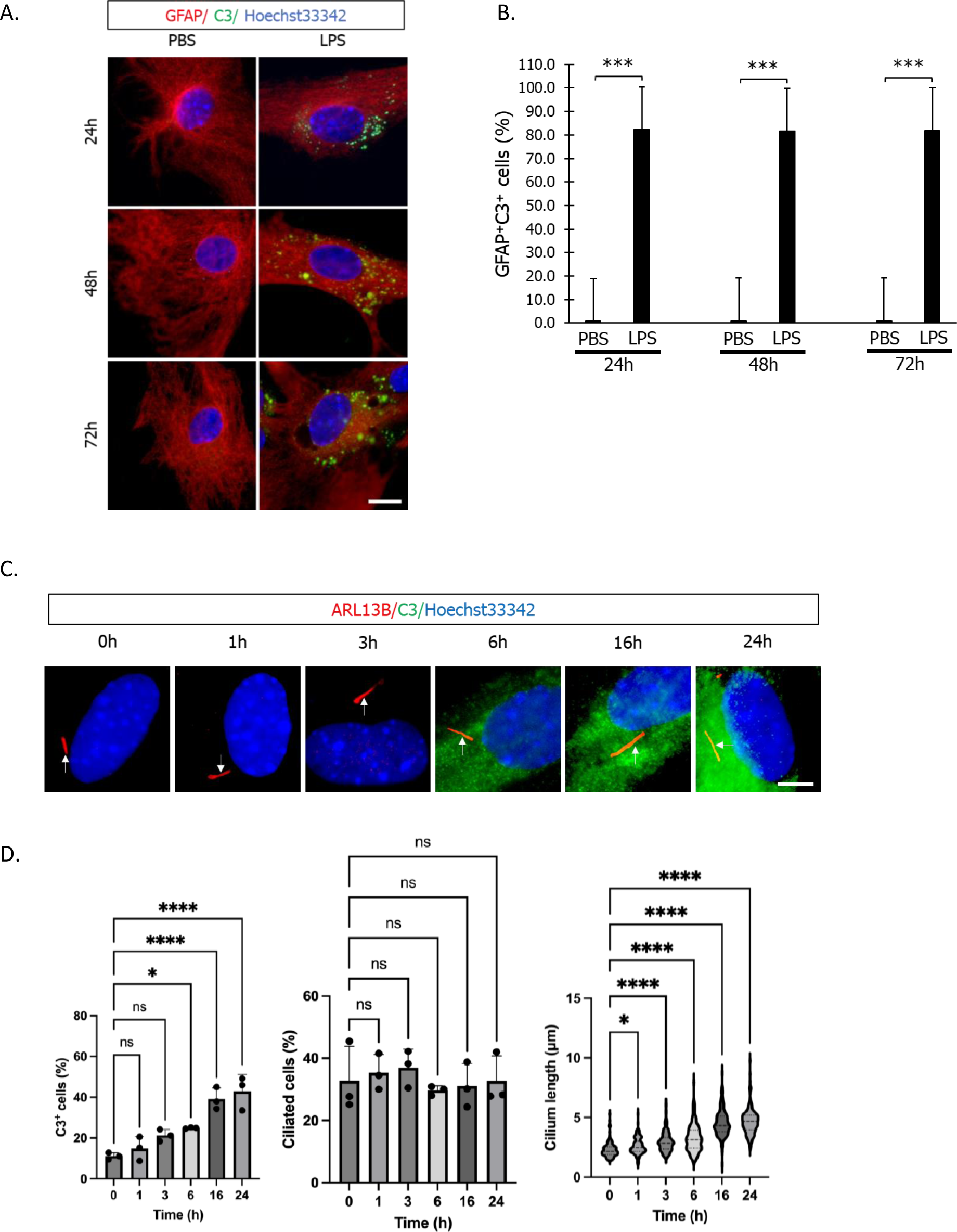
A1 astrocyte differentiation elongates primary cilium length. (A) Representative immunostaining image of C3 (green) and GFAP (astrocyte marker, red). Mixed cortical glial cells were treated with 100 ng/ml LPS for 24 h, 48 h, and 72 h. Scale bar; 20 μm. (B) Astrocytic primary cilium length shown in Figure S1A. Three independent experiments were performed. ***, P <0.001 (Mann‒Whitney U test). (C) Temporally-dependent C3 expression levels and cilium length were increased astrocyte cultures treated with 3 ng/mL IL-1⍺, 30 ng/ml TNF-⍺, and 400 ng/ml C1q. Representative images of Arl13B (red), C3 (green) and nuclei (Hoechst 33342, blue) are shown. Three independent experiments were performed. Scale bar; 5 μm. (D) The percentage of C3-positive cells (left) and ciliated cells (middle) and cilium length (right) shown in Figure S1C. Three independent experiments were performed. *, P <0.05, ****, P <0.0001, ns, nonsignificant. (Tukey’s multiple comparison).

**Figure S2.**
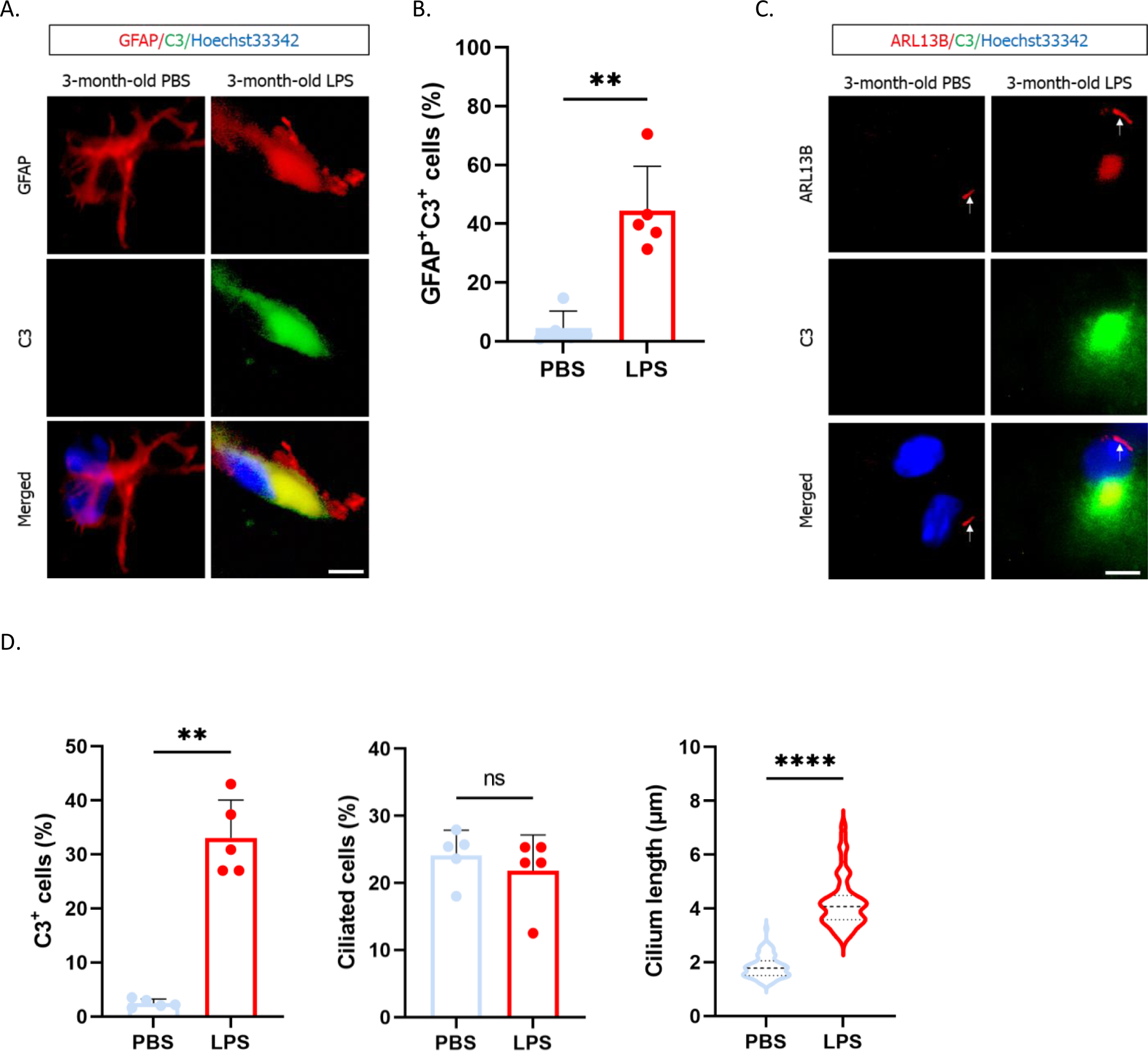
LPS injection into mice elongates the astrocytic primary cilium and induces cognitive decline. (A) Representative images of GFAP (red), C3 (green), and nuclei (blue) in the mouse brain. Mice were i.p. injected with 1 mg/kg LPS twice a week for 6 weeks. PBS-treated mice; n=5. LPS-treated mice; n=5. Scale bar: 5 µm. (B) Percentage of GFAP-positive, C3-positive cells shown in Figure S2A. **, P <0.01 (Mann‒Whitney U test). (C) Representative images of Arl13B (red), C3 (green), and nuclei (blue) in the mouse brain. PBS-treated mice; n=5. LPS-treated mice; n=5. Scale bar: 5 µm. (D) Percentage of C3-positive cells (left), percentage of ciliated cells (middle), and cilium length (right) shown in Figure S2C. **, P <0.01, ****, P <0.0001, ns, nonsignificant. (Mann‒Whitney U test).

**Figure S3.**
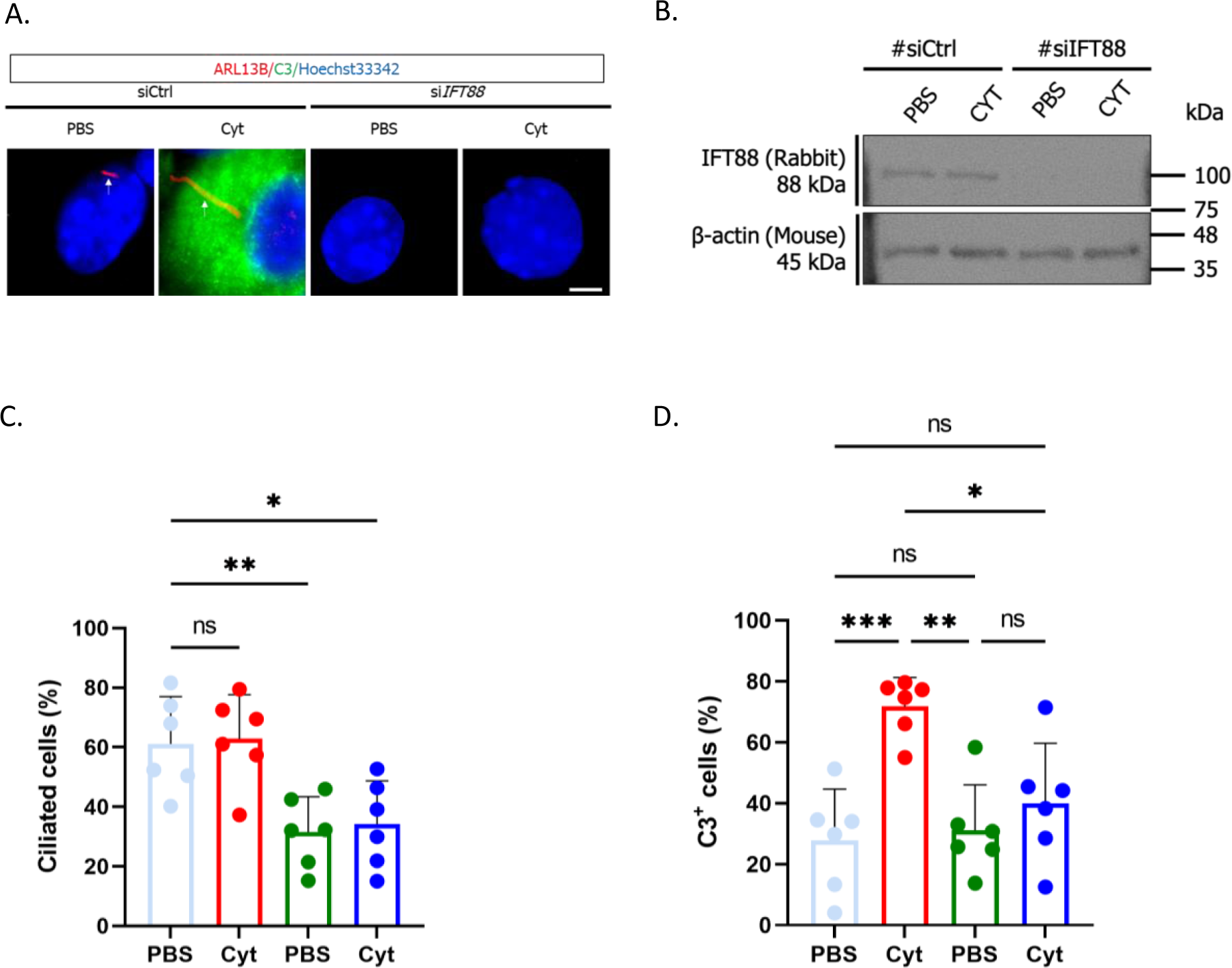
The transient downregulation of *IFT88* expression in astrocytes reduces the expression levels of C3. (A) Representative image of Arl13B (red) and C3 (green) in primary enriched astrocytes transfected with siRNA targeting *IFT88* (si*IFT88*). Nontarget siRNA was used as a control (siCtrl). Two days after transfection, cells were stimulated with 3 ng/mL IL-1α, 30 ng/ml TNFα, and 400 ng/ml C1q (Cyt) for 24 h. PBS was used as a negative control. Scale bar; 2 µm. n=6 independent experiments. (B) Representative image of IFT88 expression detected by western blotting. n=6 independent experiments. (C) Percentage of Arl13-positive primary ciliated cells shown in Figure S3A. *, P <0.05, **, P <0.01, ns, nonsignificant. (Tukey’s multiple comparison). (D) Percentage of C3-positive cells shown in Figure S3A. *, P <0.05, **, P <0.01, ***, P <0.001, ns, nonsignificant. (Tukey’s multiple comparison).

**Figure S4.**
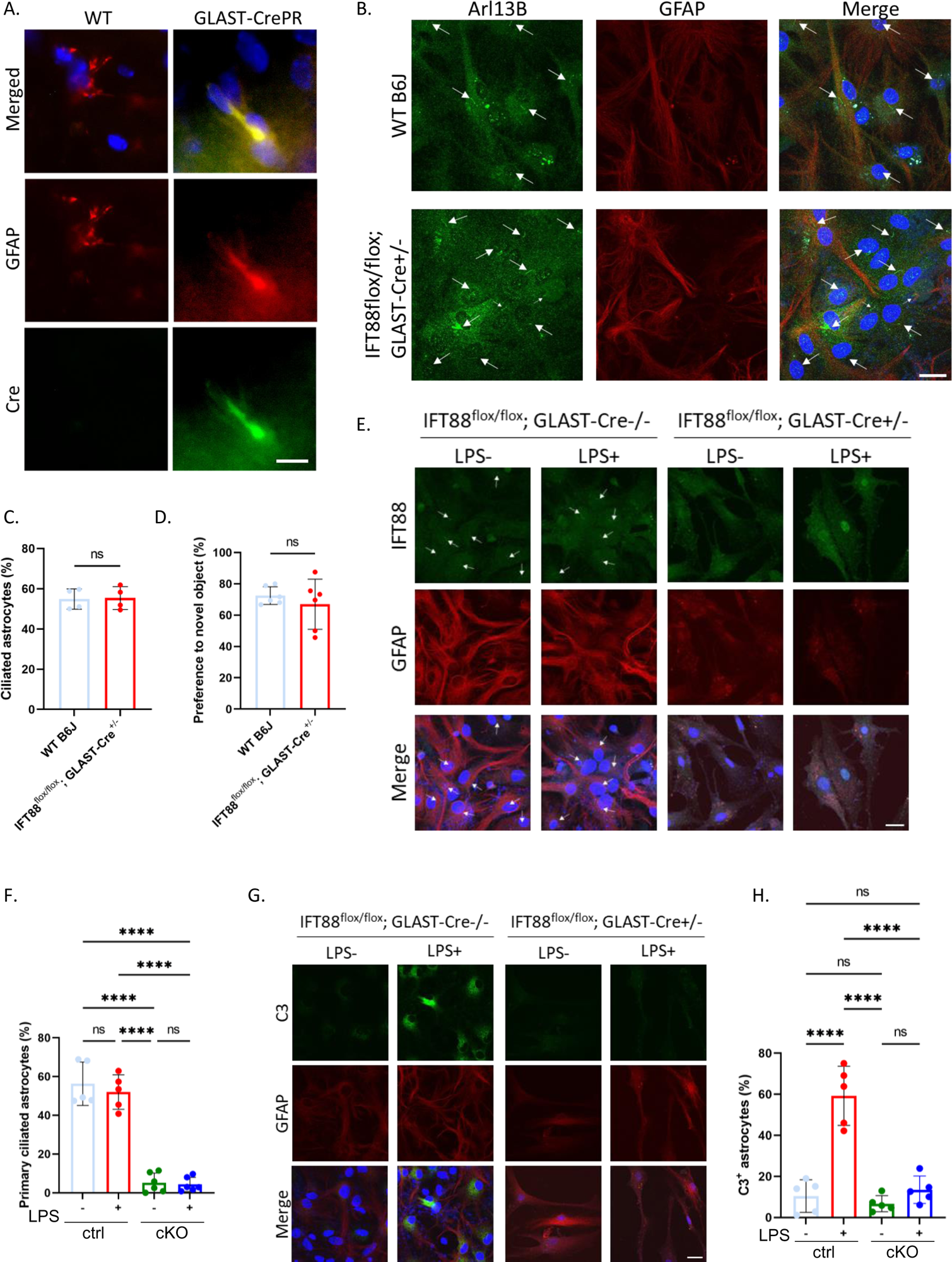
Astrocyte-specific *IFT88* gene knockout downregulates C3 expression induced by LPS stimulation. (A) Representative image of Cre recombinase expression in the mouse brain. Mouse brains were immunostained with GFAP (red), Cre (green), and Hoechst 33342 (blue). WT; C57BL/6J mice, n=3. GLAST-CrePR; *GLAST-CrePR*^+/-^ mice, n=3. Scale bar: 10 µm. (B) Representative image of Arl13B (green), GFAP (red), and Hoechst 33342 (blue) in primary astrocytes. Glial mixture cultures were prepared from C57BL/6J (WT B6J) or *IFT88*^flox/flox^; *GLAST-CrePR*^+/-^ P7 pups. The arrow indicates the primary cilium. Scale bar; 20 µm. (C) The percentage of ciliated GFAP-positive astrocytes shown Figure S4B shows in the graph. ns, nonsignificant. (Mann‒Whitney U test). n=4 independent experiments. (D) Results of behavioural (object recognition) studies in 3-month-old C57BL/6J (WT, n=6) and *IFT88*^flox/flox^; *GLAST-CrePR*^+/-^ mice (n=6). ns, nonsignificant. (Mann‒Whitney U test). (E) Representative image of IFT88 (green), GFAP (red) and Hoechst 33342 (blue) in glial mixture culture. Glial mixture cultures were prepared from P7 pups and stimulated with 10 nM RU486 for 48 h. After stimulation with RU486, cells were stimulated with 100 ng/ml LPS for 24 h. The arrow indicates the primary cilium. Scale bar; 20 µm. *IFT88*^flox/flox^; *GLAST-CrePR*^-/-^ (ctrl) pups (n=5), *IFT88*^flox/flox^; *GLAST-CrePR*^+/-^ (cKO) pups (n=6). (F) The percentage of ciliated GFAP-positive astrocytes shown Figure S4E shows in the graph. ****, P <0.0001, ns, nonsignificant (Tukey’s multiple comparison). (G) Representative image of C3 (green), GFAP (red) and Hoechst 33342 (blue) in glial mixture culture. Glial mixture cultures were prepared from P7 pups and stimulated with 10 nM RU486 for 48 h. After stimulation with RU486, cells were stimulated with 100 ng/ml LPS for 24 h. Scale bar; 20 µm. *IFT88*^flox/flox^; *GLAST-CrePR*^-/-^ (ctrl) pups (n=5), *IFT88*^flox/flox^; *GLAST-CrePR*^+/-^ (cKO) pups (n=5). (H) The percentage of C3-positive astrocytes shown Figure S4G shows in the graph. ****, P <0.0001, ns, nonsignificant (Tukey’s multiple comparison).

